# Neighbor QTL: an interval mapping method for quantitative trait loci underlying plant neighborhood effects

**DOI:** 10.1101/2020.05.20.089474

**Authors:** Yasuhiro Sato, Kazuya Takeda, Atsushi J. Nagano

## Abstract

Phenotypes of sessile organisms, such as plants, rely not only on their own genotype but also on the genotypes of neighboring individuals. Previously, we incorporated such neighbor effects into a single-marker regression using the Ising model of ferromagnetism. However, little is known about how to incorporate neighbor effects in quantitative trait locus (QTL) mapping. In this study, we propose a new method for interval QTL mapping of neighbor effects, named “Neighbor QTL”. The algorithm of neighbor QTL involves the following: (i) obtaining conditional self-genotype probabilities with recombination fraction between flanking markers, (ii) calculating neighbor genotypic identity using the self-genotype probabilities, and (iii) estimating additive and dominance deviation for neighbor effects. Our simulation using F2 and backcross lines showed that the power to detect neighbor effects increased as the effective range became smaller. The neighbor QTL was applied to insect herbivory on Col × Kas recombinant inbred lines of *Arabidopsis thaliana*. Consistent with previous evidence, the pilot experiment detected a self QTL effect on the herbivory at *GLABRA1* locus. We also observed a weak QTL on chromosome 4 regarding neighbor effects on the herbivory. The neighbor QTL method is available as an R package (https://cran.r-project.org/package=rNeighborQTL), providing a novel tool to investigate neighbor effects in QTL studies.

## 1 Introduction

Sessile organisms, such as land plants, have no active mobility to escape neighboring individuals. Field studies have shown that the phenotypes of an individual plant depend not only on their own genotype but also on those of neighboring plants (Barbosa et al., 2009). Such neighbor effects are mediated by direct interactions (e.g., competition and volatile communication) or indirect interactions (e.g., herbivore and pollinator movements), and thus modulate complex traits throughout a plant life cycle, including growth (Subrahmaniam et al., 2018), defense (Schuman et al., 2015; Sato, 2018; Tamura et al., 2020), and reproduction (Underwood et al., 2020). There is increasing appreciation that plant-plant interactions within a species may result in increased yield and population-wide pest resistance (Zeller et al., 2012; Wuest and Niklaus, 2018; Yang et al., 2019). However, knowledge remains limited about how to analyze the quantitative trait locus (QTL) underlying plant neighborhood effects.

QTL mapping is a well-established approach to analyze the loci responsible for complex traits (Broman et al., 2003; Broman and Sen, 2009; Broman et al., 2019). Although genome-wide association studies (GWAS) have now been developed, there are several limitations of this approach such as false positive signals due to the population structure (Hayes, 2013) and small-effect variants being overlooked if they are rare in the sample population (Korte and Farlow, 2013). While recombination events are limited in experimental crosses, the experimental approaches would overcome the problem of population structure and rare variants. In plant genetics, once GWAS leads us to find a pair of target accessions, its biparental population is then subject to QTL mapping (Sonah et al., 2015; Han et al., 2018). Therefore, QTL mapping provides a complementary analysis for GWAS to further dissect complex traits in plant genetics and breeding (Sonah et al., 2015; Rishmawi et al., 2017; Han et al., 2018; Marchadier et al., 2019).

Using the Ising model of statistical physics, our previous study proposed “Neighbor GWAS” that combined neighbor effects and a linear mixed model (Sato et al., 2019b). The core idea of neighbor GWAS was to consider the Ising model as an inverse problem of single-marker regression and, thereby, estimate the effects of neighbor genotypic identity on a trait. However, QTL mapping of neighbor effects is more complicated than single-marker analysis because QTL studies employ the maximum likelihood method for interval mapping between flanking markers (Haley and Knott, 1992; Jansen, 1993; Broman and Sen, 2009). Such an interval mapping requires a stepwise inference from genotype imputation to phenotype prediction. First, conditional genotype probabilities are obtained from the observed marker genotypes and recombination fractions between flanking markers. Then, phenotypes are inferred using the conditional genotype probabilities and marker effects (Haley and Knott, 1992). To adopt interval mapping for neighbor effects, it is necessary to define the effects of neighbor genotypic identity on a quantitative trait.

In this study, we developed an interval mapping method for testing neighbor effects in QTL studies. The proposed method, “Neighbor QTL”, was applied to simulated data and recombinant inbred lines (RILs) of *Arabidopsis thaliana*. Furthermore, the new QTL method was built into an R package.

## 2 Materials and Methods

### 2.1 Model

We first developed a basic regression model and then defined QTL effects for interval mapping. To combine neighbor effects and a linear model, we focused on the well-known model of statistical physics, Ising model (McCoy and Maillard, 2012). The Ising model defines magnetic energy arising from physical interactions among neighboring magnets. By analogy, we regarded an individual as a magnet, genotypes as dipoles, and a trait as energy. Given the observed traits (or energy), we estimated interaction coefficients of the Ising model to infer neighbor effects.

#### 2.1.1 Joint regression for self and neighbor effects

To incorporate neighbor effects into a linear regression, we developed a joint model following the single-marker regression of neighbor GWAS (Sato et al., 2019b). We considered a situation where a number of inbred lines occupied finite sites in a two-dimensional space and assumed that an individual is represented by a magnet, whereby two homozygotes at each marker, AA or BB, correspond to north or south dipole (Fig. 1). We defined *x_i_* or *x_j_* as the genotype at a focal marker respectively for *i*-th focal individual or *j*-th neighbor, where *x_i_*_(*j*)_ ∈{AA, BB} = {1, −1}. We then used multiple regression to model the effects of self-genotype and neighbor genotypic identity on a trait of *i*-th individual *y_i_* as

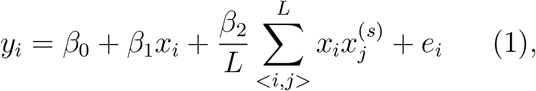

where *β*_0_, *β*_1_, and *β*_2_ indicated intercept, self-genotype, and neighbor effects, respectively. The residual for a trait value of the focal individual *i* was denoted as *e_i_*. The neighbor covariate 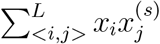 was the sum of products for all combinations between the *i*-th focal individual and the *j*-th neighbor at the *s*-th scale of spatial distance from the focal individual *i* (Fig. 1). The total number of neighbors *L* varied in response to the spatial scale *s* to be referred. The coefficient of neighbor effects *β*_2_ was scaled by *L*. If two individuals shared the same genotype at a given locus, the product *x_i_x_j_* became positive; the product *x_i_x_j_* became negative if two individuals had different genotypes. Thus, the effects of neighbor genotypic identity on a trait *y_i_* was dependent on the coefficient *β*_2_ and the number of two genotypes in a neighborhood.

**Figure 1:**
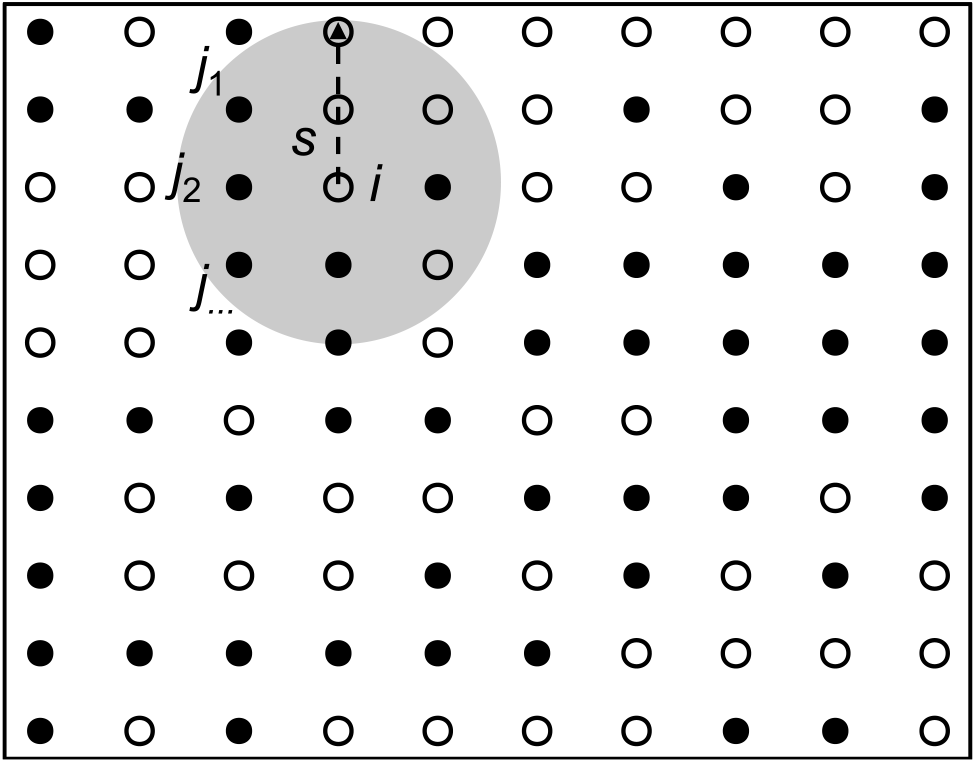
Assumption of neighbor effects in a two-dimensional space. A white or black point indicates an individual having AA or BB genotype, respectively. A grey circle shows an effective area of neighbor effects at the spatial distance *s* from the focal individual *i*. Neighbor effects then occur depending on genotype similarity between the focal individual *i* and all the *j*-th neighbors within the spatial distance *s*.

Notably, the multiple regression model eq. 1 was posed as an inverse problem of the Ising model. When summing up *y_i_* for all individuals and substituting coefficients as *E* = −*β*_2_*/L*, *H* = −*β*_1_ and *ϵ_I_* = ∑(*y_i_* − *β*_0_), eq. 1 could be transformed into the total magnetic energy of a two-dimensional Ising model as 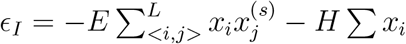 (McCoy and Maillard, 2012). In such a case, the neighbor effect *β*_2_ and self-genotype effect *β*_1_ could be interpreted as the interaction coefficient *E* and external magnetic force *H*, respectively.

#### 2.1.2 QTL effects of neighbor genotypic identity

To exchange a linear regression into a QTL model, we defined QTL effects for self and neighbor effects. With heterozygosity incorporated, we redefined *x_i_* and *x_j_* by a marker genotype for an *i*-th focal individual and *j*-th neighbor as *g_i_* and *g_j_*, respectively. Self QTL effects expected from those genotypes were denoted as *g_i_*_(*j*)_ ∈{AA, AB, BB} = {*a*, *d*, −*a*}, where *a* and *d* indicated additive and dominance deviation, respectively. Assuming two possible directions, we then defined QTL effects by neighbor genotypic identity between the individual *i* and *j* (Table 1). Given the QTL effects of self and neighbor effects, we decomposed a trait of *i*-th individual *y_i_* as

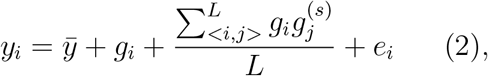

where 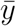 and *e_i_* indicated a population mean of traits and a residual for the focal individual *i*, respectively. Assuming that two marker effects *a* and *d* were unlikely to be equivalent between self and neighbor effects, we introduced *a*_1_ and *d*_1_ to the self QTL effects; and *a*_2_ and *d*_2_ to the neighbor QTL effects. If QTL effects were completely additive (i.e., *a*_1_ = *a*_2_ = 1 and *d*_1_ = *d*_2_ = 0), the QTL model eq. 2 had the same structure as the linear regression eq. 1. In such an additive model, the coefficients *β*_1_ and 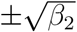 represented additive QTL effects. It was also worth noting that the sign of 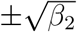 determined the direction of the effects of genotypic identity on a trait (Table 1).

**Table 1:**
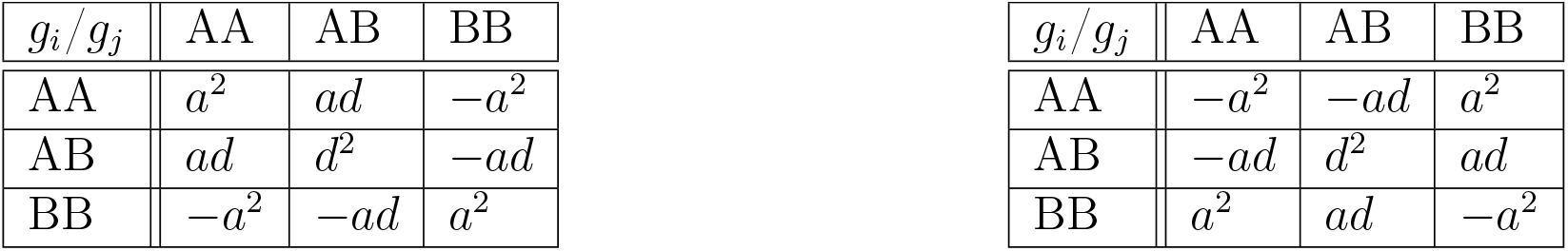
QTL effects expected by genotypic identity between the individuals *i* and *j* with AA, AB, or BB genotypes. The additive and dominance deviation is represented by *a* and *d*, respectively. The left table shows a case in which a share of same QTL genotypes exerts positive effects on a trait *y_i_*, whereas the right table shows a case in which a share of same genotypes exerts negative effects on *y_i_*

#### 2.1.3 Interval mapping for neighbor genotypic identity

To enable interval mapping, we extended the single-marker QTL model eq. 2 to multiple pseudo-markers. In particular, we modified Haley-Knott regression that approximated the maximum likelihood method by a simple regression (Haley and Knott, 1992; Broman and Sen, 2009). The proposed algorithm consisted of three steps: (i) obtaining conditional self-genotype probabilities, (ii) calculating neighbor genotypic identity from the conditional self-genotype probabilities, and (iii) regressing trait values on the conditional self-genotype probabilities and neighbor genotypic identity.

The first step to obtain conditional self-genotype probabilities was the same as that of standard QTL mapping. Let *p_i_*_(*j*)_ be the probability for the focal individual *i* or neighbor *j* to have a certain genotype at an interval pseudo-marker. We defined the conditional self-genotype probability for the individual *i* as *p_i_* = Pr(*g_i_* = {AA, AB, BB}|**M**), and obtained *p_i_* from the number of observed markers × *n* individuals matrix **M** and its recombination fraction following hidden Markov models (Lander and Green, 1987; Broman et al., 2003). Based on the products of the conditional self-genotype probabilities, we further calculated the conditional probabilities for neighbor genotypic identity *p_i_p_j_*. We then defined *g_i_g_j_* as the QTL effects by neighbor genotypic identity; and *p_i_p_j_* as the expected probability for two genotypes to interact, whereby the expected neighbor QTL effects was *p_i_p_j_g_i_g_j_*. These probabilities were summed up for all possible combinations of the genotypes as 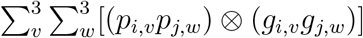, where the subscript *v* and *w* indicated the three genotype states AA, AB, and BB.

Similar to Haley-Knott regression, we finally estimated the QTL effects *g_i_* and *g_i_g_j_* by regressing the trait values *y_i_* on *p_i_* and 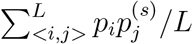, respectively. The additive and dominance deviation for the self QTL effects *a*_1_ and *d*_1_ were considered as average differences in trait values among AA, AB, or BB genotypes, such that 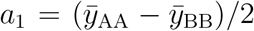 and 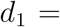 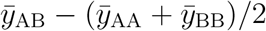 (Broman and Sen, 2009). In such a case, the regression coefficient *β*_1_ gave 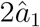 when −1, 0, and 1 dummy groups were assigned for the AA, AB, and BB genotypes, respectively, or gave 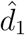 when 0, 1, and 0 were assigned for the AA, AB, and BB genotypes, respectively (Broman et al., 2003).

For neighbor QTL effects, the additive and dominance deviation *a*_2_ and *d*_2_ were also considered as the average differences in trait values among the nine possible combinations (Table 1) as 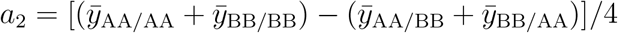 and 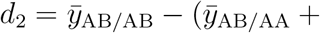 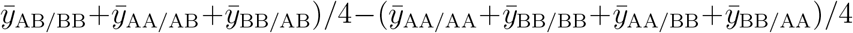. In this case, trait values *y_i_* could be fitted by a quadratic regression on the group of nine genotype combinations (Fig. 2). Suppose that 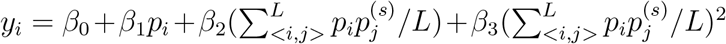 represents such a quadratic regression, where the linear or quadratic coefficient *β*_2_ or *β*_3_ provides estimates for the additive or dominance deviation 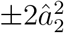 or 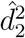, respectively. Practically, we could estimate *a*_2_ and *d*_2_ by the quadratic regression of the trait values *y_i_* on the neighbor genotypic identity 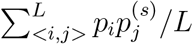, with nine genotype combinations encoded as AA/AA, BB/BB, AA/AB, AB/AA, AB/AB, AB/BB, BB/AB, AA/BB, BB/AA = {1, 1, 0.25, 0.25, 0.0, −0.25, −0.25, −1, −1}.

**Figure 2:**
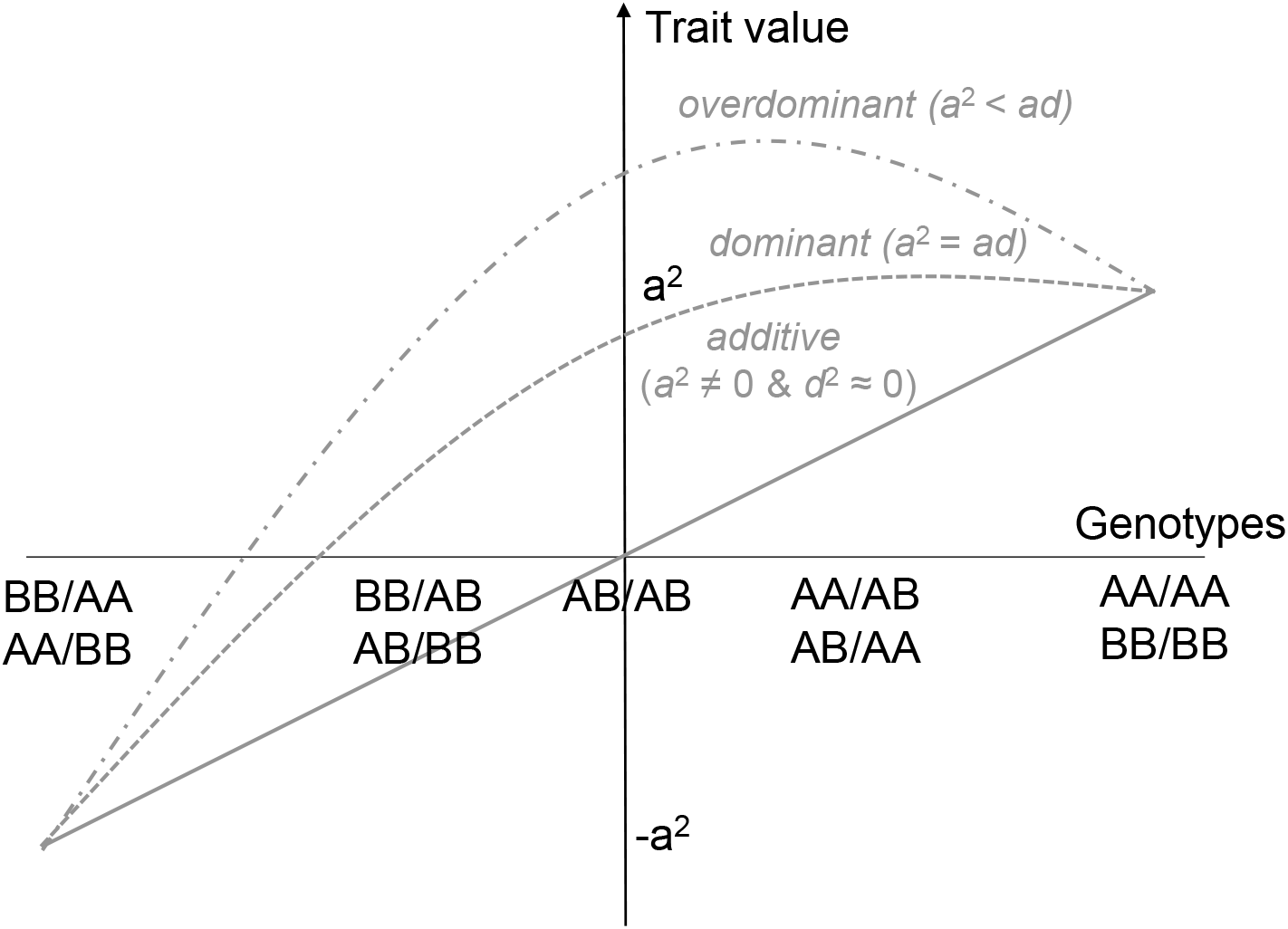
A scheme explaining approximation of neighbor QTL effects by quadratic regression. Trait values *y_i_* are regressed on nine possible combinations of genotype identity between a focal individual *i* and its neighbor *j* (Table 1). The additive or dominance deviation *a* or *d* is represented by the linear or quadratic term, respectively. If the linear coefficient is negative, it indicates the case in which AA/AA and BB/BB combinations had negative QTL effects on traits (Right of Table 1).

Based on the linear and quadratic regression, we decomposed a trait *y_i_* into self and neighbor QTL effects. To distinguish the two effects, we estimated *a*_1_, *d*_1_, *a*_2_, and *d*_2_ by following the six-step iterations.

1. Estimate *a*_1_ by a linear regression on self-genotype probabilities, with −1, 0, and 1 encoded for the AA, AB and BB genotypes, respectively.
2. Estimate *d*_1_ by a linear on self-genotype probabilities regression, with 0, 1, and 0 encoded for the AA, AB and BB genotypes, respectively.
3. Calculate self QTL effects with 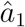 and 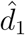.
4. Include the self QTL effect as a covariate at a focal marker.
5. Estimate ±*a*_2_ and *d*_2_ by a quadratic regression on neighbor genotypic identity, with [−1, 1] dummy groups assigned for nine genotype combinations.
6. Calculate joint QTL effects with 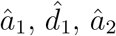 and 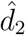.

Based on 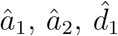 and 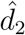, we inferred 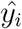 and derived log_e_-likelihood (LL) from model deviance. LOD score for the self or neighbor effects were designated as LOD_self_ = log_10_[exp(LL_self_ − LL_null_)] or LOD_nei_ = log_10_[exp(LL_nei_ − LL_self_)], which could be obtained in steps 3 and 6, respectively.

When there were only two genotypes, the quadratic regression was replaced by a linear regression to estimate the additive neighbor effects. For the case of inbred lines lacking AB heterozygotes, we estimated the additive deviation *a*_2_ by a linear regression of trait values *y_i_* on the neighbor genotypic identity 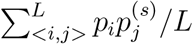, with 1 and −1 dummy groups assigned for the AA and BB genotypes, respectively. In case of backcross lines lacking BB homozygotes, the additive deviation corresponded to the dominance deviation so that *d*_2_ = −*a*_2_. The additive deviation *a*_2_ could be estimated by a linear regression with the AA and AB genotypes encoded into −1 and 0, respectively. These two linear models were equivalent in the sense that both inbred and backcross lines had two genotypes with additive effects.

#### 2.1.4 Variation partitioning with the QTL model

Prior to the genome scan, we estimated the effective spatial scale *s* by calculating the proportion of phenotypic variation explained (PVE) by neighbor effects. Incorporating two random effects into a linear mixed model, we were able to partition phenotypic variation into PVE by self effects, neighbor effects, and residuals (Sato et al., 2019b). According to previous studies (Henderson et al., 1959; Kang et al., 2008), the linear mixed model was expressed as

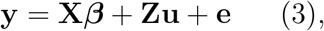

where **y** indicated a phenotype vector as *y_i_* ∈ **y**; **X*β*** indicated fixed effects with a matrix including a unit vector and all covariates **X** and a coefficient vector ***β***; **Zu** indicated random effects with *u_i_* ∈ **u** and a design matrix **Z**; and **e** indicates residuals where *e_i_* ∈ **e**. The random effects and residuals were further decomposed as 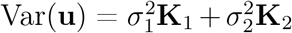 and 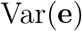 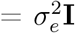, where the *n* × *n* individuals similarity matrix for self-genotype or neighbor identity was scaled by the number of markers *q* as 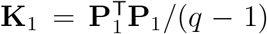 or 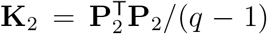, respectively. Given that one of two alleles is similar between heterozygotes and homozygotes, here we defined additive polygenic effects for self QTLs as *g_i_* ∈{AA, AB, BB} = {−1, 0, 1}; and for neighbor QTLs as *g_i_g_j_* ∈{AA/AA, BB/BB, AA/AB, AB/AA, AB/AB, AB/BB, BB/AB, AA/BB, BB/AA} = {1, 1, 0.5, 0.5, 0.0, −0.5, −0.5, −1, −1}. In these cases, the *q* × *n* matrix **P**_1_ included expected self-genotype values as elements 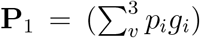 and **K**_1_ represented a kinship matrix that was calculated from all the pseudo-markers (Broman et al., 2019). Similarly, the *q* × *n* matrix **P**_2_ included the neighbor genotypic identities as elements 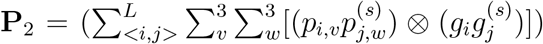 and **K**_2_ represented a genome-wide structure of neighbor genotypic identity. Based on the three variance component parameters, we calculated PVE by self or neighbor effects as 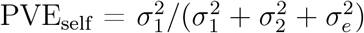 or 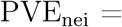 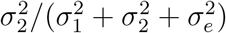. Additionally, the heritability was designated as 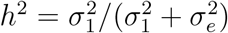 when 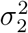 was set at 0.

Using the linear mixed model eq. 3, our previous simulations revealed that the effective spatial scale of neighbor effects could be determined by increasing the patterns of PVE_nei_ from *s* = 0 to a large *s* (Sato et al., 2019b). If the effective range was narrow, PVE_nei_ approached to a plateau at a small value of *s*. In contrast, PVE_nei_ linearly increased with *s* if the effective range was broad. To generalize these results for a continuous two-dimensional space, here we introduced ΔPVE metric as differences in PVE from *s* to *s* + 1 such that ΔPVE = PVE_nei,*s*+1_ − PVE_nei,*s*_. Using such differential metrics, we quantified how PVE_nei_ approached to a plateau across *s* as follows:

1. Categorize spatial scales as *s* ∈ *S* based on the percentiles for pairwise Euclidean distance between individuals.
2. Calculate PVE_nei_ from *s* = 1 to the maximum elements of *S*.
3. Calculate ΔPVE_nei_ and determine *s* = arg max ΔPVE_nei_

The proposed algorithm using a differential PVE was called “ΔPVE method” hereafter.

#### 2.1.5 An R package, “rNeighborQTL”

In addition, the neighbor QTL method was built into an R package, “rNeighborQTL”. The rNeighborQTL took as input objects from the R/qtl package (Broman et al., 2003), allowing us to save phenotypes and genotypes as common “cross” objects. Because of the stepwise testing, the self QTL effects yielded the same results as standard QTL mapping. For the ΔPVE method, the mixed models eq. 3 were solved using the algorithm of average information restricted maximum likelihood (AI-REML) (Gilmour et al., 1995) implemented in the gaston package (Perdry and Dandine-Roulland, 2020). An additional, but necessary, input file was a spatial map describing the positions of individuals at the x- and y-axes. The rNeighborQTL package is available via CRAN at https://cran.r-project.org/package=rNeighborQTL.

The rNeighborQTL package included several options to analyze a variety of QTL data. Alternative to linear (mixed) models (eq. 1 and eq. 3), logistic (mixed) models could also be selected to handle a binary phenotype (Faraway, 2016; Chen et al., 2016). Because the logistic mixed model did not provide 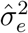 (Chen et al., 2016; Perdry and Dandine-Roulland, 2020), PVE_nei_ was substituted by the ratio of phenotypic variation explained (RVE) by neighbor effects as 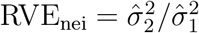, when a binary trait was subject to the ΔPVE method. The neighbor QTL also allowed additional covariates when conducting a genome scan. This option enabled composite interval mapping (Jansen, 1993), if genetic markers other than a focal locus were considered covariates. When a significant marker was detected by the single-QTL analysis, it was also possible to test two-way interactions, such as namely epistasis, between the neighbor QTL effects across a genome. Details are documented in the rNeighborQTL package.

### 2.2 Simulation

We performed a benchmark test using simulated data on F2 and backcross lines. With a random spatial map generated, we simulated neighbor effects based on “fake.f2” and “fake.bc” autosome genotypes implemented in the R/qtl package (Broman et al., 2003). The spatial positions were sampled from a uniform distribution Unif(1, 100) across a continuous two-dimensional space. We estimated *a*_1_ for self-phenotypes of “fake.f2” and “fake.bc” data after the trait values were scaled to have a mean of zero and variance of 1, and assigned 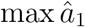 to a randomly selected marker. In contrast to the major-effect marker, small coefficients, i.e., 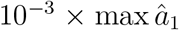, were assigned to the other markers to simulate polygenic effects. Additive (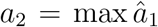 and 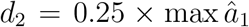), dominant 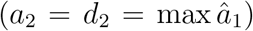, and overdominant (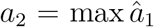 and 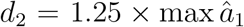) scenarios were analyzed for the F2 lines, while only additive scenario (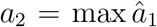 and 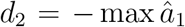) was applicable for the backcross lines. Thirty traits were simulated for true effective distances given at ten to fifty percentiles of pairwise Euclidean distance among individuals. The simulated neighbor effects were added to the self-phenotypes of “fake.f2” or “fake.bc” dataset, with 75% of phenotypic variation being attributable to the neighbor effects. Then we applied the ΔPVE method and a genome scan for the joint traits, and calculated LOD_nei_ at *s* = arg max ΔPVE_nei_ to evaluate the power to detect neighbor effects.

### 2.3 Data

To apply the neighbor QTL on real data, we conducted a pilot QTL experiment using the yellow-striped flea beetle *Phyllotreta striolata* and RILs of *Arabidopsis thaliana* (Fig. S1A). Adults of flea beetles access host plants by jumping, and leaf holes made by these beetles are easily countable. These flea beetles are known to prefer glabrous *A. thaliana* to hairy accessions (Sato et al., 2019a). To observe large phenotypic variation in leaf holes, we selected RILs derived from hairy and glabrous accessions in this study.

#### 2.3.1 Plants and insects

We used 130 accessions, including parental and recombinant inbred lines between Col(*gl1*) and Kas-1 accession (Wilson et al., 2001). Col(*gl1*) plants produce no trichomes, while Kas has sparse trichomes on leaves and stems. The RILs are known to vary in the trichome production, disease resistance (Wilson et al., 2001), and flowering time (Li et al., 2006). The genotype data were available in Wilson et al. (2001). The set of RILs was obtained through the Arabidopsis Biological Resource Center (ABRC) (Stock ID, CS84999: https://abrc.osu.edu/).

Flea beetles were maintained under a long-day condition (16:8 hours light:dark cycles with a 22 °C constant air temperature) in an environmental chamber (Biotron LH-241PFDSP, NK system, Osaka, Japan). To establish the experimental population, we collected ca. 200 adults from *Brassica* cultivars grown in the field within Otsu City, Shiga Prefecture, Japan (35°01′N 135°51′E) during November 2018 and May 2019. Adults of *P. striolata* consume shoots and especially prefer to young glabrous leaves, whereas larvae consume below-ground tissue of *Brassica* plants; therefore, we reared adults and larvae on leaves and swollen hypocotyls, respectively. Young leaves of Boc choy *Brassica rapa* var. *chinensis* or Chinese cabbage *B. rapa* var. *pekinensis* were supplied for the adults. The larvae were allowed to feed on swollen hypocotyls of the radish *Raphanus sativus* var. *longipinnatus* or the turnip *B. rapa* subsp. *rapa* buried in moisten vermiculite. Adult females laid eggs in the moisten vermiculite, and it took a month (28 to 32 days) for eggs to become adults.

#### 2.3.2 Experimental procedure

To investigate neighbor effects in herbivory, we allowed adult beetles to feed on RIL seedlings grown in a plastic cell tray. Three seeds for each accession were sown on each compartment of the cell tray (13 × 10 cells composed of 20 × 20 mm^2^ compartment) with the accessions randomized. The seeds were acclimated under a constant dark condition with 4 °C for seven days, and then allowed to germinate under a long-day condition (16:8 hours light:dark cycles with a 20 °C constant air temperature). The seedlings were grown under the long-day condition for 24 days, with 2000-fold diluted liquid fertilizer (N:P:K = 6:10:5; Hyponex, Hyponex Japan, Osaka) supplied once. On day 14 after the germination, the seedlings were thinned out to leave one seedling per compartment. Prior to the feeding experiment, we recorded the presence or absence of leaf trichomes and the occurrence of bolting by direct observation and determined the rosette diameter (mm) by analyzing seedling images using Image J software (Abràmoff et al., 2004). The cell tray was enclosed by a white mesh cage (length 29.2 cm × width 41.0 cm × height 27.0 cm: Fig. S1B). Thirty adult beetles were released into the cage and allowed to feed on plants for 72 hours. We counted leaf holes as a measure of herbivory for each plant as flea beetles left small holes when they fed on leaves (Fig. S1C). The final sample size was 126 individuals; out of 130 accessions, 4 accessions (CS84877, CS84873, CS84950, and CS84894) were not germinated, CS84898 lacked genotype data, and CS84958 had two replicates of individuals. The data are included in the rNeighborQTL package.

#### 2.3.3 Data analysis

We used R version 3.6.0 (R Core Team, 2019) for all statistical analyses. A genetic map for the Col × Kas RILs was estimated using the est.map() function in the R/qtl package (Broman et al., 2003). Self-genotype probabilities were calculated using the calc.genoprob() function implemented in the R/qtl package (Broman et al., 2003). The number of leaf holes was log-transformed and analyzed using linear models. The presence of trichomes and bolting was analyzed using logistic models. When analyzing the number of leaf holes, we incorporated the presence or absence of bolting, the rosette diameter, and the edge (or not) of the cell tray into covariates. The neighbor QTL was performed using the rNeighborQTL package developed above. Examples using the Col × Kas dataset are available in the vignette of rNeighborQTL package, where the usage of each function is also documented. A genomewide significance level was determined by empirical percentiles of the maximum LOD score among 999 permuted traits. We considered *p* < 0.1 and *p* < 0.05 a suggestive and significant level, respectively. We also set an arbitrary threshold at LOD score of 1.5 when discussing the results.

## 3 Results and Discussion

### 3.1 Simulation using F2 and backcross lines

We simulated neighbor effects based on “fake.f2” and “fake.bc” data implemented in the R/qtl package (Broman et al., 2003). The maximum additive deviation of self QTL effects, 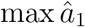, was 0.56 and 0.28 for F2 and backcross lines, respectively. These values were assigned for neighbor QTL effects to achieve similar a signal strength between self and neighbor effects, while minor effects were allocated to other loci. Considering the polygenic variation as random effects, we applied the ΔPVE method for simulated traits. The estimated distance given by *s* = arg max ΔPVE increased as the true distance increased (Fig. 3), indicating that the ΔPVE method was effective.

**Figure 3:**
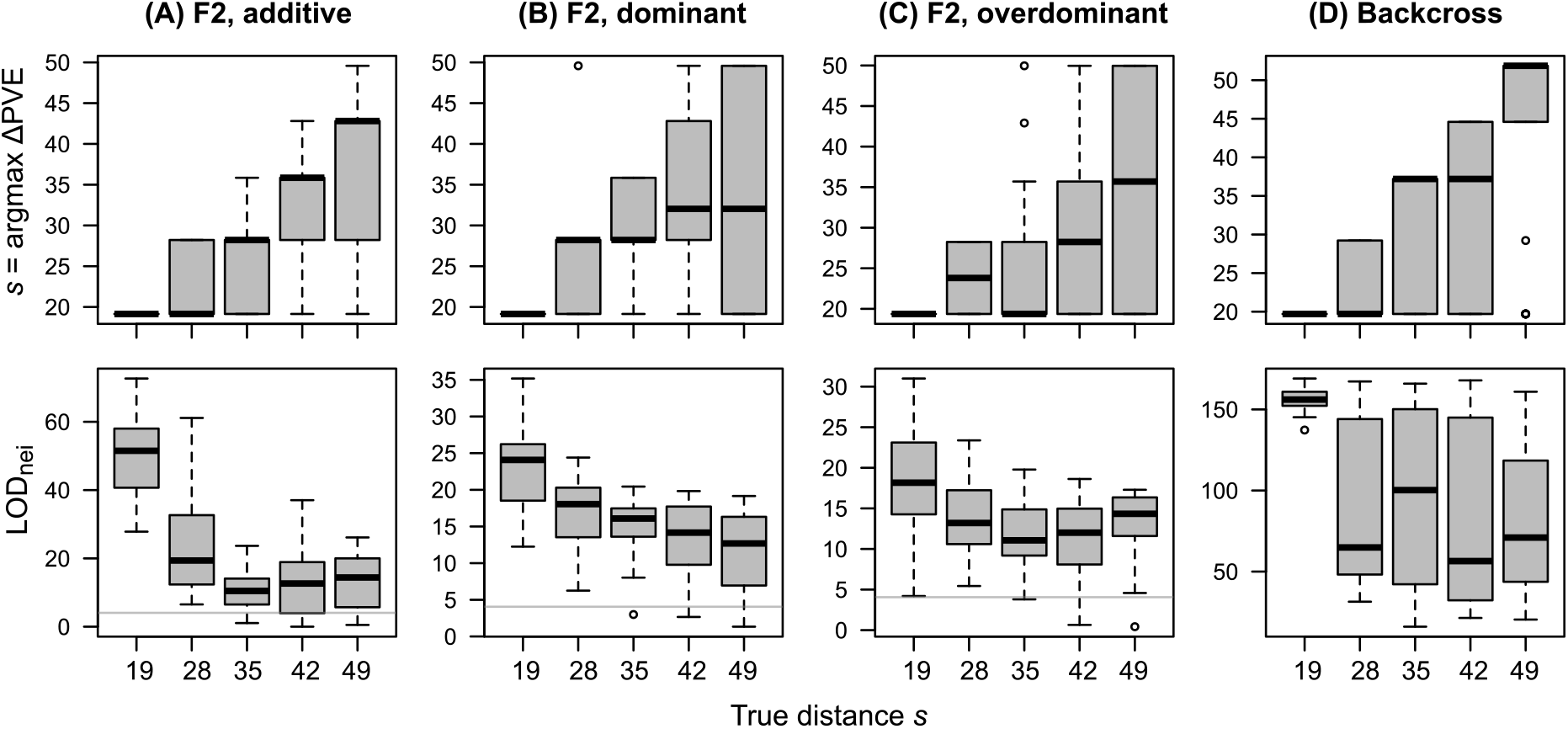
Benchmark test using simulated F2 and backcross datasets. Upper panels show the distance estimated by ΔPVE method, while lower panels show LOD_nei_ of a major-effect marker at the estimated distance. The x-axis corresponds to ten to fifty percentiles of pairwise Euclidean distance. Thirty traits were simulated for each distance class. Boxplots represent median by a center line; upper and lower quartiles by box limits; 1.5× interquartile range by whiskers; and outliers by points. Horizontal lines indicate a LOD threshold at *p* = 0.05 after Bonferroni correction.

When the efficient distance of neighbor effects was limited, such short-range neighbor effects were well detected using ΔPVE method and the quadratic approximation (median LOD_nei_ > 4 at the ten percentile of pairwise Euclidean distance: Fig. 3). Although the power to detect long-range neighbor effects was lowered, LOD score was still larger than the Bonferroni threshold (median LOD_nei_ > 4: Fig. 3). These results indicated that short-range neighbor effects could be detected in any scenario, although it was relatively difficult to detect long-range effects.

For backcross lines, both short- and long-range neighbor effects were well detected (median LOD_nei_ > 4 for all *s*: Fig. 3D). The backcross lines had two genotypes with the additive deviation alone and were well fitted using linear approximation (Fig. 3D), whereas the additive traits for F2 lines were less likely fitted using the quadratic model assuming three genotypes with the additive and dominance deviation (Fig. 3A). Given the model structure underlying F2 lines, it is plausible that the quadratic term was unnecessary for the additive F2 traits, indeed, it decreased the power to detect neighbor effects.

### 3.2 Self QTL effects in Col × Kas RILs

The observed number of leaf holes ranged from 0 to 38 with a median of 4 (Fig. S1D). The total variation in the number of leaf holes was explained at 5% by the trichome production; 2% by bolting; 10% by the rosette diameter; and 22% by the edge effects (Analysis-of-Variance, *F* = 9.1, 3.7, 20.7, and 43.4; *p* = 0.003, 0.06, 10^−4^, and 10^−8^, respectively). With the kinship matrix **K**_1_ considered a random effect, a linear mixed model estimated the heritability as 5.6% for the leaf holes, though it was not significant (Likelihood ratio test, 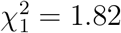, *p* = 0.18).

To scan self QTL effects, we conducted standard QTL mapping of the trichome production, the number of leaf holes, and bolting (Fig. 4; Table 2). For self QTL effects on the trichome production, we detected a strong peak near the *GLABRA1* locus (>20 LOD_self_ score: Fig. 4B). Considering the rosette diameter and bolting covariates, we observed a suggestive but the largest self QTL effect on the leaf holes at the *GLABRA1* locus (LOD_self_ = 1.97: Fig. 4C). For the bolting, we observed the largest significant peak on the bottom of chromosome 1 (>4 LOD_self_), and the second largest and suggestive peak on the top of chromosome 4 (LOD_self_ = 1.92: Fig. 4D).

**Table 2:** Estimated QTL effects in Col × Kas RILs of *Arabidopsis thaliana*. Markers with any >1.5 LOD scores (highlighted by bold letters) are shown. Additive effects 2*a*_1_ indicate the effect size when Kas alleles are replaced by two Col alleles, while 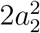 indicates the effect size of identical homozygotes over different ones. The sign of 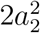 determines the direction of neighbor genotypic identity (Table 1). The LOD_nei_ score is shown on the spatial distance at which ΔPVE peaks.

| Trait | Marker | Chr | Position (cM) | $2a_1$ | LOD<sub>self</sub> | $\pm 2a_2^2$ | LOD<sub>nei</sub> | Distance |
| --- | --- | --- | --- | --- | --- | --- | --- | --- |
| Trichome | GL1 | 3 | 65.24 | **-2.83** | **22.8** | 3.28 | 0.13 | 7.82 |
| Holes | GL1 | 3 | 65.24 | **0.21** | **1.97** | -0.25 | 0.05 | 7 |
|  | nga8 | 4 | 0 | -0.07 | 0.13 | **2.67** | **1.86** | 7 |
| Bolting | nga692 | 1 | 102.0 | **-1.04** | **4.15** | -0.72 | 0.13 | 2.24 |
|  | R30025 | 3 | 126.1 | 0.06 | 0.01 | **2.82** | **1.80** | 2.24 |
|  | nga8 | 4 | 0 | **1.05** | **1.92** | 0.41 | 0.02 | 2.24 |

**Figure 4:**
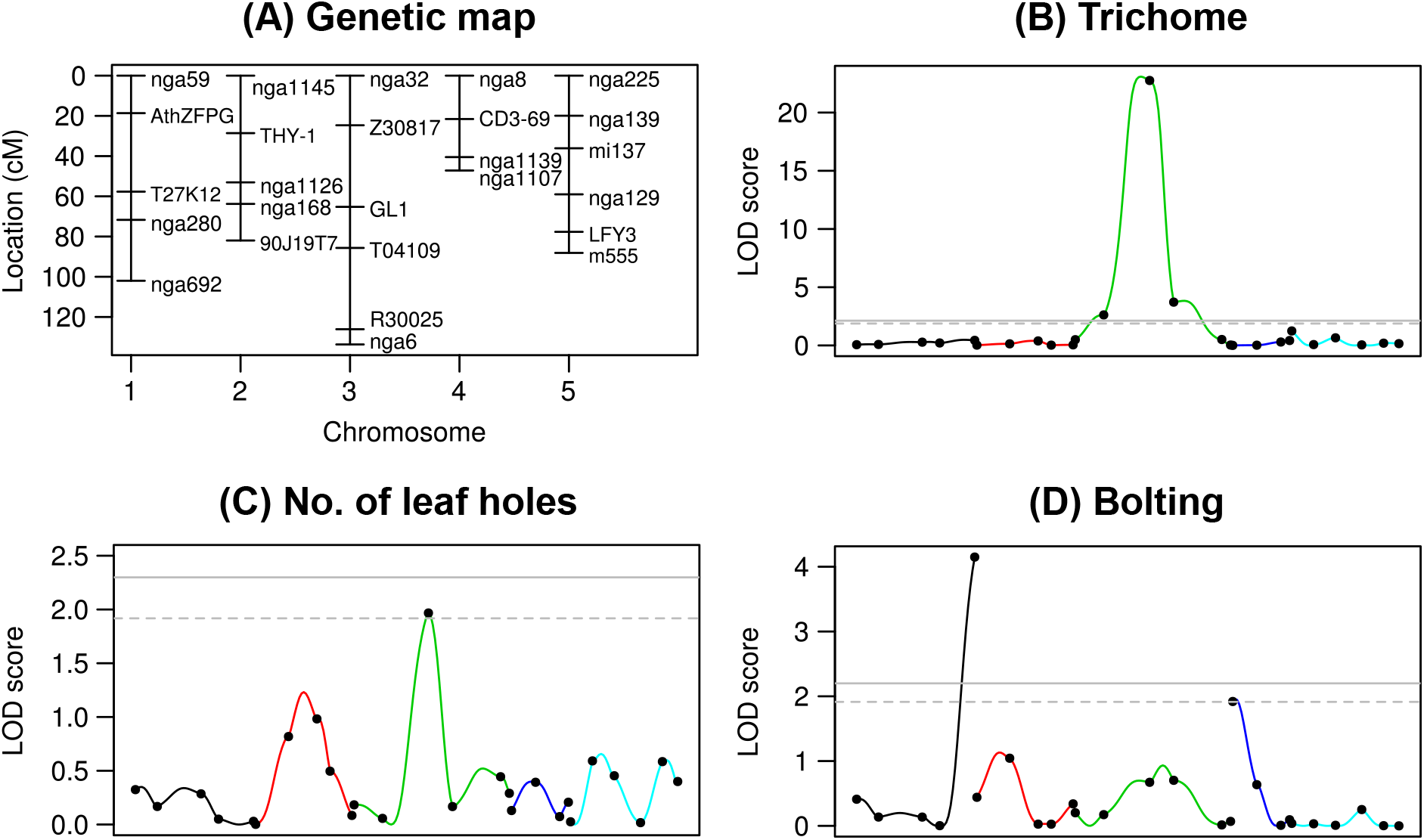
Genetic map and LOD scores for self QTL effects in Col × Kas RILs. (A) Genetic map showing the locations of 26 markers among the five chromosomes of *Arabidopsis thaliana*. LOD_self_ score for the trichome production (B), the number of leaf holes (C), bolting (D). Colors correspond to chromosome numbers, and dots indicate observed markers. A solid and dashed horizontal line indicates a significant (*p* < 0.05) and suggestive (*p* < 0.1) LOD threshold with 999 permutations, respectively.

Several studies reported the same QTLs or a particular gene function for the self effects on trichomes, defense, and flowering. Remarkably, *GLABRA1* gene on the chromosome 3 is known to encode a myb transcription factor regulating leaf trichome developments (Ishida et al., 2008) and deter feeding by flea beetles (Sato et al., 2019a). The result that *GLABRA1* possessed self QTL effects on the leaf holes adds a biological value to the insect herbivory data. Furthermore, two self-bolting QTLs on the chromosome 1 and 4 were located near flowering time QTLs in Col × Kas RILs (Li et al., 2006). Thus, our pilot experiment supports previous evidence for the loci responsible for plant development and defense.

### 3.3 Neighbor QTL effects in Col × Kas RILs

To estimate the effective distance of neighbor effects, we applied ΔPVE method with every ten percentile categories for pairwise Euclidean distance (Fig. 5). For the number of leaf holes, the ΔPVE_nei_ was peaked at 7 distance scale from a focal individual (Fig. 5A), covering almost all the experimental arena from the center plant. At this estimated distance, the neighbor effects explained 6% of total variation in the leaf holes, with the both self and neighbor similarity **K**_1_ and **K**_2_ considered random effects in linear mixed models. At 7.8 distance scale, the neighbor effects explained 8.7% of total variation in leaf holes at the maximum, though it was not significant compared to its heritability by self QTL effects (Likelihood ratio test, 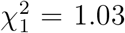, *p* = 0.31). It seemed plausible that the effective distance was relatively large for insect herbivory, because the adult beetles were likely free to move within the small experimental cage. On the other hand, the ΔPVE became the largest at the nearest scale for the bolting and explained over a half variation compared to the self QTL effects (RVE_nei_ = 0.68 at *s* = 2.24: Fig. 5B), suggesting that the bolting was unlikely affected by distant neighbors. For the trichome production, ΔPVE method revealed that there were little variation explained by neighbor effects (RVE_nei_ ≈ 0 otherwise models failed to converge).

**Figure 5:**
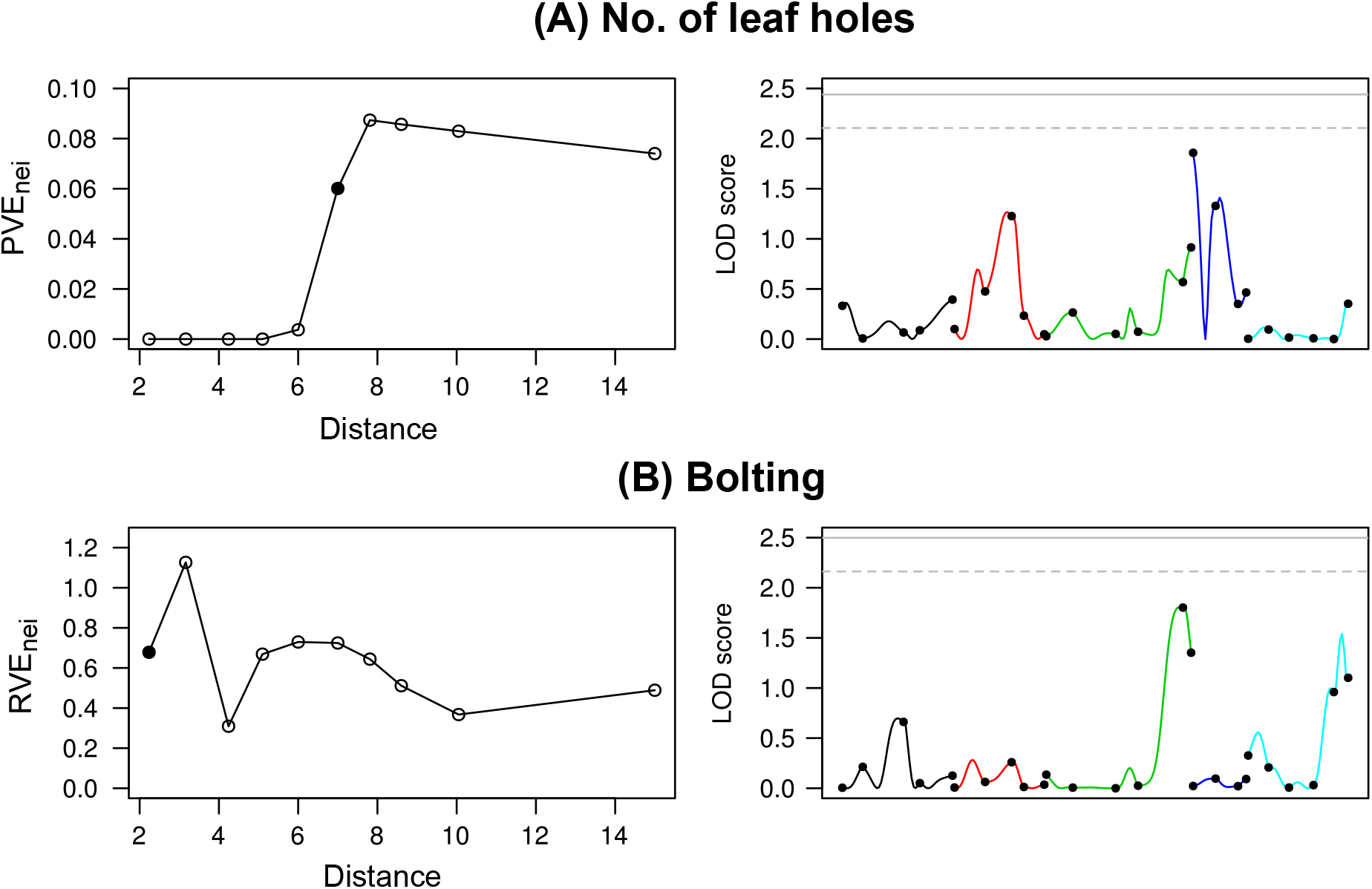
Phenotypic variation explained and LOD score attributed to neighbor effects on the number of leaf holes (A) or the presence of bolting (B) in Col × Kas RILs. Left: Proportion or ratio of phenotypic variation explained by neighbor effects (PVE_nei_ or RVE_nei_) plotted against the pairwise distance among individuals. A closed point indicates the distance at which ΔPVE peaked. Right: LOD_nei_ score for neighbor QTL effects at the distance at which ΔPVE peaked. Colors correspond to chromosome numbers, and dots indicate observed markers. A solid and dashed horizontal line indicates a significant (*p* < 0.05) and suggestive (*p* < 0.1) LOD threshold with 999 permutations, respectively.

A genome scan for neighbor effects was performed using the estimated spatial distance (Fig. 5; Table 2). Regarding the neighbor QTL effects on the leaf holes, we observed, although weak, the largest QTL on the top of chromosome 4 at the nga8 marker (LOD_nei_ = 1.86), which was also the position the second largest self-bolting QTL occurred. This neighbor QTL had no significant epistasis as shown by < 1.1 LOD score for all the two-way interactions between the nga8 and other markers (Fig. S2A). Neither the neighbor QTL nor *GLABRA1* locus detected above was overlapped with known self-QTLs of powdery mildew resistance (Wilson et al., 2001), suggesting independence of the herbivory QTLs on the disease resistance loci. At the nearest scale for bolting, we found a weak neighbor QTL at the R30025 marker on the chromosome 3 (LOD_nei_ = 1.8: Fig. 5; Table 2). This QTL did not have any significant epistasis with the other markers (< 1.1 LOD score: Fig. S2B).

Ecological studies have shown that how easy an individual plant is to find, termed plant apparency, drives neighbor effects through visual crypsis against herbivores (Hambäck et al., 2000; Castagneyrol et al., 2013; Strauss et al., 2015). In the present study, the neighbor QTL involved in the leaf holes was located near a self-bolting QTL at the top of chromosome 4, suggesting the potential importance of plant apparency in neighbor effects in anti-herbivore defense. In addition, the positive sign of the additive neighbor effects *a*_2_ at that marker indicated that the number of leaf holes decreased when neighbors had different genotypes (Table 2). This implies that the mixture of flowering and vegetative plants may acquire population-wide resistance to flea beetles since the effective distance of neighbor effect was sufficiently large to encompass almost the entire experimental arena. These results led us to hypothesize that the self QTL underlying plant apparency might facilitate population-wide anti-herbivore defense, called associational resistance (Hambäck et al., 2000), through its pleiotropy on neighbor effects.

### 3.4 Further applicability and limitation

Theoretical advantage of the Ising model lies in its inference of spatial arrangements that optimize total magnetic energy. Once the self and neighbor coefficients are estimated by the marker-based regression, these two coefficients may infer which genotype distributions can minimize or maximize the population-sum of trait values (Sato et al., 2019b). In the context of neighbor QTL, additive effects suggest that positive and negative *a*_2_ favors clustered or mixed patterns for maximizing the sum of trait values, respectively. However, in cases where dominance effects and epistasis are involved, how such a complex genetic basis affects the optimal spatial arrangement remains unexplored. These potential effects of genetic architecture on a population-level outcome of neighbor effects would be of theoretical as well as empirical interest for future studies.

Superior to the previous neighbor GWAS, the present neighbor QTL has a flexibility to deal with heterozygosity. However, the use of neighbor QTL is still restricted to autosomes because sex-dependent inheritance of neighbor effects remains unknown. Standard QTL mapping on sex chromosomes is known to require from one to three degree-of-freedom (Broman et al., 2006), and thus its extension to neighbor effects may be more complex than the self QTL effects. In addition, the neighbor QTL approximated the maximum likelihood method by a quadratic regression, in which phenotype variance was assumed to be equal among the nine combinations among three QTL genotypes. Our simulation revealed that the quadratic approximation could handle the overdominance, but became inferior to linear approximation if additive effects alone governed a trait. We should thus be aware of statistical models behind the neighbor QTL. Practically, both the intercross and the inbred models might be utilized if a sample population is partially inbred.

### 3.5 Conclusion

The present neighbor QTL, together with the previous neighbor GWAS (Sato et al., 2019b), provides a sort of novel tools to incorporate neighbor effects into quantitative genetics. These methods may provide insights into genetic architecture underlying neighbor effects as exemplified by the pilot study of insect herbivory on *A. thaliana*. Once the neighbor GWAS screens candidate accessions, their crossed progeny can be inspected by the neighbor QTL. The line of R packages, “rNeighborQTL” and “rNeighborGWAS”, would help investigate neighbor effects using a complementary set of GWAS and QTL data.

## Acknowledgements

The authors are grateful to E. Yamamoto for helpful comments on the draft; to Dynacom Co., Ltd. for technical assistance on the R package development; and to L. G. Kawaguchi for collecting insect materials from the field. This study was supported by Japan Science and Technology Agency (JST) PRESTO (Grant number, JPMJPR17Q4) and Japan Society for the Promotion of Science (20K15880) to Y.S., and JST CREST (JPMJCR15O2) to A.J.N. No conflicts of interests concern this study.

## Supplementary Materials

**Figure S1:**
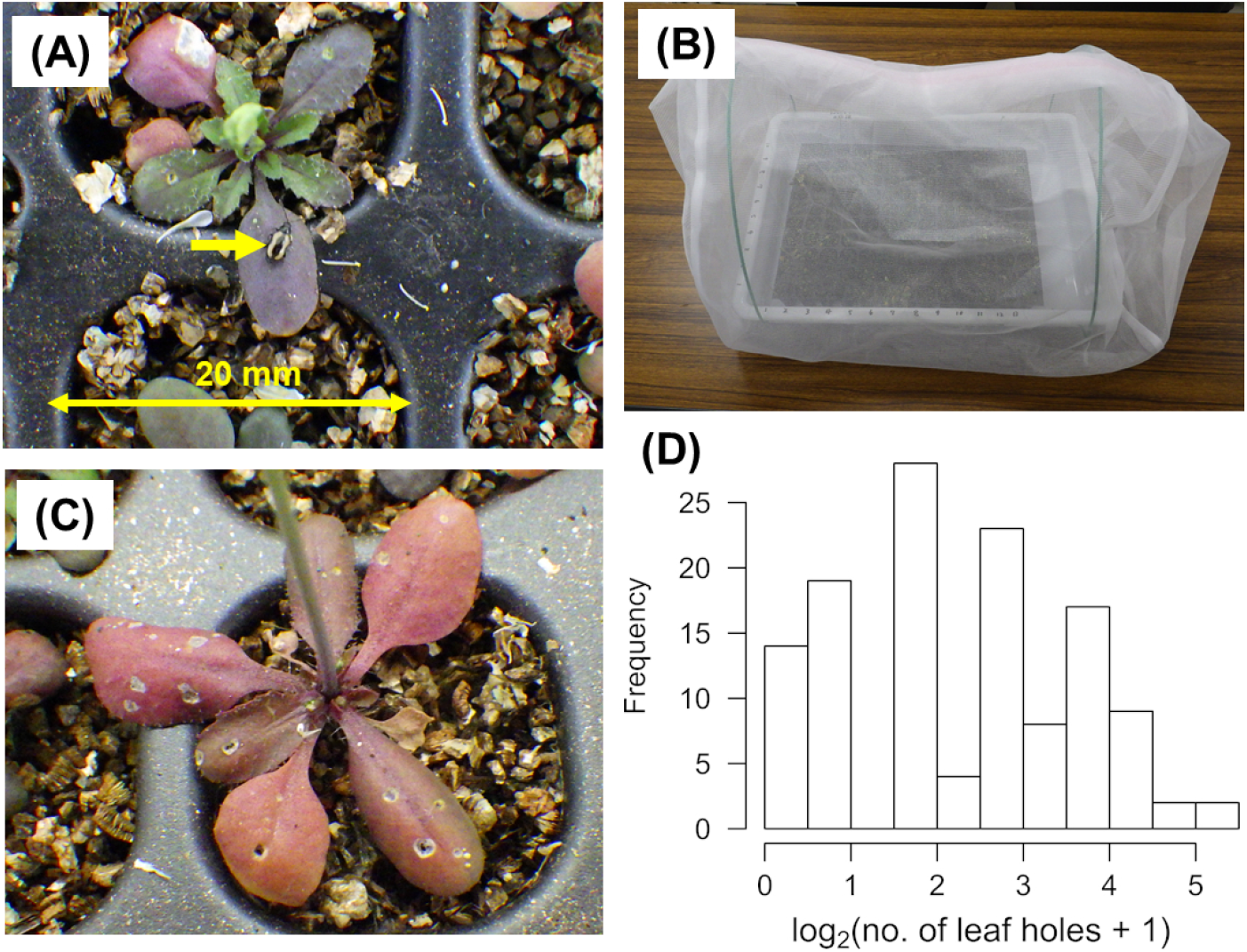
Pilot QTL experiment using *Arabidopsis thaliana* and the yellow-striped flea beetle *Phyllotreta striolata*. (A) An adult beetle on a vegetative plant. (B) An experimental cage including 130 Col × Kas RILs. (C) A plant attacked by *P. striolata*. Leaf holes were made by adult beetles. (D) Histogram for the observed number of leaf holes.

**Figure S2:**
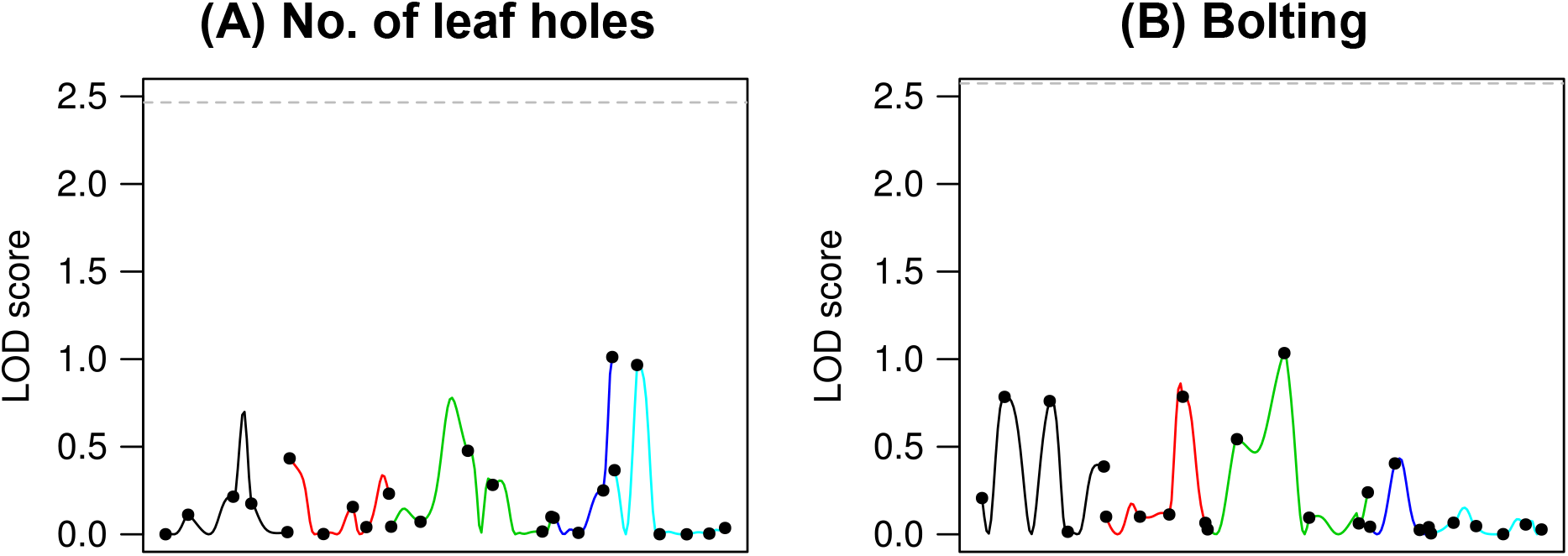
Epistasis in neighbor QTL effects on the number of leaf holes between the nga8 and other markers (A); or on the presence of bolting between the R30025 and other markers (B). Colors correspond to chromosome numbers, and dots indicate observed markers. A dashed horizontal line indicates a suggestive (*p* < 0.1) LOD threshold with 999 permutations.

## Notes

https://cran.r-project.org/package=rNeighborQTL

### Competing Interest Statement

The authors have declared no competing interest.

### Summary of Updates

Abstract revised; Introduction revised; Figure 1 added

